# *De novo* acquisition of antibiotic resistance in six species of bacteria

**DOI:** 10.1101/2024.07.03.601945

**Authors:** Xinyu Wang, Alphonse de Koster, Belinda B. Koenders, Martijs Jonker, Stanley Brul, Benno H. ter Kuile

**Affiliations:** Biology and Microbial Food Safety, Swammerdam Institute for Life Sciences, University of Amsterdam, Amsterdam, The Netherlands; RNA Biology & Applied Bioinformatics, Swammerdam Institute for Life Sciences, University of Amsterdam, Amsterdam, The Netherlands

## Abstract

Bacteria can become resistant to antibiotics in two ways, by acquiring resistance genes through horizontal gene transfer and by *de novo* development of resistance upon exposure to non-lethal concentrations. The importance of the second process, *de novo* build-up, has not been investigated systematically over a range of species and may be underestimated as a result. To investigate the DNA mutation patterns accompanying the *de novo* antibiotic resistance acquisition process, six bacterial species encountered in the food chain were exposed to step-wise increasing sublethal concentrations of six antibiotics to develop high levels of resistance. Phenotypic and mutational landscapes were constructed based on whole genome sequencing (WGS) sequencing at two time points of the evolutionary trajectory. In this study, we found: 1) all of the six strains can develop high levels of resistance against most antibiotics. 2) increased resistance is accompanied by different mutations for each bacterium-antibiotic combination. 3) the number of mutations varies widely, with *Y. enterocolitica* having by far the most. 4) in the case of fluoroquinolone resistance, a mutational pattern of *gyrA* combined with *parC* is conserved in five of six species. 5) mutations in genes coding for efflux pumps are widely encountered in gram-negative species. The overall conclusion is that very similar phenotypic outcomes are instigated by very different genetic changes.

**IMPORTANCE:** The significance of this study lies in the comparison of how six species of distinct genomic background under uniform conditions develop high levels of antibiotic resistance against six antibiotics. The mutational patterns in these six species of bacteria identify common target mutations and reveal how they acquire mutations from various pathways to survive and grow when exposed to sub-lethal levels of antibiotics. In addition to providing insights in microbial genetics, outcome of this study will assist policymakers when formulating practical strategies to prevent development of antimicrobial resistance in human and veterinary health care.

## INTRODUCTION

Exposure to non-lethal concentrations of antimicrobials causes bacteria to develop de novo resistance (1, 2). In agriculture low concentrations of antimicrobials have been and are sometimes still being applied as growth promoters and to prevent infectious diseases (3). The rapid build-up of resistance in *E. coli* as a consequence of exposure to step-wise increasing concentrations of antibiotics has been documented extensively (4–7). The steady increase of resistance levels in microbes isolated from livestock and foodstuffs is well documented (8, 9). The contribution of de novo selected resistance is not known quantitatively but might be considerable. The resistance that eventually is transferred to human health care hampers treatment of infectious diseases and increases the costs of healthcare (10).

Five resistance mechanisms can be highlighted for bacteria to combat antibioticś action: reduced permeability, antibiotic efflux pumps, modifying the target molecule or target pathways, inactivation of the drug and adaptation of affected pathways to bypass a blocked step (11, 12). Whole genome sequencing (WGS) analysis comparing *E. coli* strains made resistant to the original naïve strain identified DNA mutations that accompany acquisition of resistance (6) and the mechanisms that enable the cell to develop resistance (5). In some, but not all, cases a clear target mutation can be identified in response to fluoroquinolones (7, 13, 14) (15) and rifampicin(16) Most studies of de novo resistance were performed on single species with the exception of the comparison of the clinically relevant ESKAPE species made resistant to the five most applied antibiotics in human health care (17). Still several questions remain to be answered: 1) Can all bacteria develop high levels of resistance against all or most antibiotics, as observed in *E. coli*? 2) Does each antibiotic trigger similar responses in different species? 3) Does any species have a consistent response to all six antibiotics? This study addresses these questions by comparing development of resistance to six different antibiotics in six species relevant for agriculture, *Salmonella enterica subsp*. *houtenae, Yersinia enterocolitica*, *Staphylococcus aureus*, *Enterococcus faecalis*, *Bacillus. subtilis* and *Acinetobacter pittii*.

## MATERIALS AND METHODS

### Bacteria and reagents

The following species were used throughout the study: *Salmonella enterica subsp*. *houtenae, Yersinia enterocolitica*, *Staphylococcus aureus*, *Enterococcus faecalis*, *Bacillus. subtilis* and *Acinetobacter pittii.* These are either species frequently encountered on livestock, or close relatives of such species, chosen because they are less virulent and thus safer to handle. Wild type *S. enterica* DSM 9221, *S. aureus* DSM 110565, *E. faecalis* ATCC 4707711, and *Y. enterocolitica* ATCC 9610 were purchased from companies DSM and ATCC. *A. pittii* is a clinical isolation and was identified by WGS sequencing(18), *Bacillus subtilis* B168 is a lab collection (19). All experiments were performed in Tryptic soy broth (TSB). Bacteria from -80 °C glycerol stock was steaked on Tryptic soy broth agar plates (TSA). The strains of *Y. enterocolitica* and *A. pittii* were cultured at 30 °C and the other four strains were cultured at 37 °C in aerobic condition to have a sufficient growth. Each of the antibiotic solutions was created from powder stock (Sigma, Germany) and was filtered using a 0.22 µm pore size (Merk, America). The antibiotic stocks were stored at -20 °C until needed. Amoxicillin was prepared freshly every four days. Antibiotics were kept in the fridge at below 4 °C for a maximum of four days.

### Antimicrobial susceptibility testing

The minimal inhibitory concentration (MIC) assays were performed with 2-fold increasing concentrations of antibiotics in TSB(4). All measurements from a single colony of the WT or evolved populations were performed with two technical replicates in 96 well plate using the Thermo Scientific Multiskan FC with SkanIt software reader at a wavelength of 595 nm. The end volume of each well was filled with 150 µl TSB with a starting 0.05 OD_595_ of overnight culture, and all measurements took 24 hours at 37 °C/30 °C. The cut-off in OD for resistance was 0.2 and the growth curve was used to monitor the validity of the MIC measurement.

### Inducing antibiotic resistance

The development of resistance in the evolution experiments was started from a single colony of each of the six WT strains. Bacteria were isolated from TSA and cultured in 5 ml TSB in tubes in a shaker incubator at 220 rpm shaking for 24 hours in one cycle. The evolution experiments were performed in stepwise increasing sublethal concentrations of six antibiotics following the standard protocol (4, 14). One strain grown in the absence of antibiotics was used as biological control. Strains were initially exposed to 1/8 of the MIC with inoculum size starting overnight culture OD_595_ of 0.1. Sequential doubling of the antibiotic concentration was performed when samples grew more than 75% of the growth observed in the biological control, measured by yield of OD. The evolution experiments continued until strains developed stable high levels of resistance. Samples of the evolved populations were stored at -80 °C in 30 % glycerol.

### DNA isolation and whole-genome sequencing

In total 144 samples of resistant strains, sampled at the middle and final time point of their evolutionary trajectory, six biological control samples and six wild type samples were subjected to genomic DNA sequencing. Isolation of genomic DNA was performed using the Purelink genomic DNA Kit (America). Genomic DNA libraries were generated using the NEBNext Ultra II FS DNA Library Prep kit for Illumina (New England BioLabs) in combination with NEBNext multiplex oligos for Illumina (96 Unique Dual Index Primer Pairs; New England BioLabs) according to the manufacturer’s instructions. In briefly, 500 ng genomic DNA was used as input with a fragmentation time of 5 min, aiming at an insert size distribution of 275–475 bp by following the corresponding size selection option provided in the protocol. The resulting size distribution of the libraries with indexed adapters was assessed using a 2200 TapeStation System with Agilent D1000 ScreenTapes (Agilent Technologies). The libraries were quantified on a QuantStudio 3 Real-Time PCR System (Thermo Fisher Scientific) using the NEBNext Library Quant Kit for Illumina (New England BioLabs) according to the instructions of the manufacturer. The libraries were clustered and sequenced (2 x 150 bp) on a NextSeq 550 Sequencing System (Illumina) using a NextSeq 500/550 Mid Output v2.5 kit (300 cycles) (Illumina). The sequencing coverage depth was set up aiming at 120x ∼ 170x on average.

### Detection of maximum exponential growth rate

The growth rate of resistant strains at the middle and final time point of their evolutionary trajectory was determined by growing them in a plate reader (Thermoscientific, multiskan FC). The OD measurements were started using overnight cultures of these strains diluted to OD_595_ of 0.05 in 150 µl TSB. The plates were incubated in the plate reader for 24 hours and the ODs were measured at 595 nm at 10 min intervals. The growth rates were analyzed in R studio, using a script that was downloaded from GitHub (https://github.com/Pimutje/Growthrates-in-R/releases/tag/Growthrates). The significance of the difference between in the growth rate between the resistant strains and controls was analyzed with a t-test. A p-value ≤ 0.05 was considered significant.

## RESULTS

### Experimental evolution of resistance by six bacteria against six antibiotics

To provide a baseline for the experiments documenting the evolution of resistance against six antibiotics by six species of bacteria, the minimal inhibitory concentrations (MICs) were measured of these species against three bactericidal and three bacteriostatic antibiotics (**Table 1**). As representatives for the beta-lactam antimicrobials amoxicillin was used for gram-negative species and cefepime for gram-positives. There are in total three cases of intrinsic resistance: *S. aureus* against enrofloxacin, *E. faecalis* against kanamycin, and *S. enterica* against erythromycin. With the exception of the bacteriostatic tetracycline and, to a lesser extent, chloramphenicol, the MIC values varied between species.

**TABLE 1.** Measurements of the MIC (μg/mL) of six bacterial species against the six antibiotics.

| $\mu\text{g/mL}$ | AMO/CEF | ENR | KAN | TET | ERY | CHL |
| --- | --- | --- | --- | --- | --- | --- |
| *S. enterica* | 1 | 0.0625 | 64 | 2 | 512 | 4 |
| *Y. enterocolitica* | 32 | 0.0625 | 64 | 1 | 64 | 4 |
| *A. pittii* | 32 | 0.0625 | 8 | 1 | 4 | 2 |
| *E. faecalis* | 16 | 4 | 1024 | 2 | 4 | 8 |
| *B. subtilis* | 4 | 0.125 | 4 | 8 | 0.5 | 8 |
| *S. aureus* | 16 | 16 | 64 | 2 | 0.5 | 32 |

AMO: amoxicillin, CEF: cefepime, ENR: enrofloxacin, KAN: kanamycin, TET:
tetracycline, ERY: erythromycin, CHL: chloramphenicol.

In order to assess the ability of microbes to acquire resistance against often used antimicrobials, six species of bacteria were exposed to stepwise increasing concentrations of six antibiotics (**Fig. 1**). Almost all combinations of a microbe and an antibiotic, below called drug/bug combinations, showed a rapid increase of the resistance. Notable exceptions were the combinations *S. enterica* and amoxicillin, *A. pittii* and enrofloxacin and *S. aureus* with tetracycline and chloramphenicol, that showed little increase. There were no general trends observable. In the three cases of intrinsic resistance, the strains of *S. enterica*, *E. faecalis*, and *S. aureus* developed still higher levels of resistance, showing increases in MIC of 2^3^, 2^3^, and 2^6^ to erythromycin, kanamycin and enrofloxacin, respectively. There was no bacterial species that consistently developed least or most resistance, nor was there an antibiotic that always showed either more or less de novo resistance (**Fig. 2**).

**FIG. 1.**
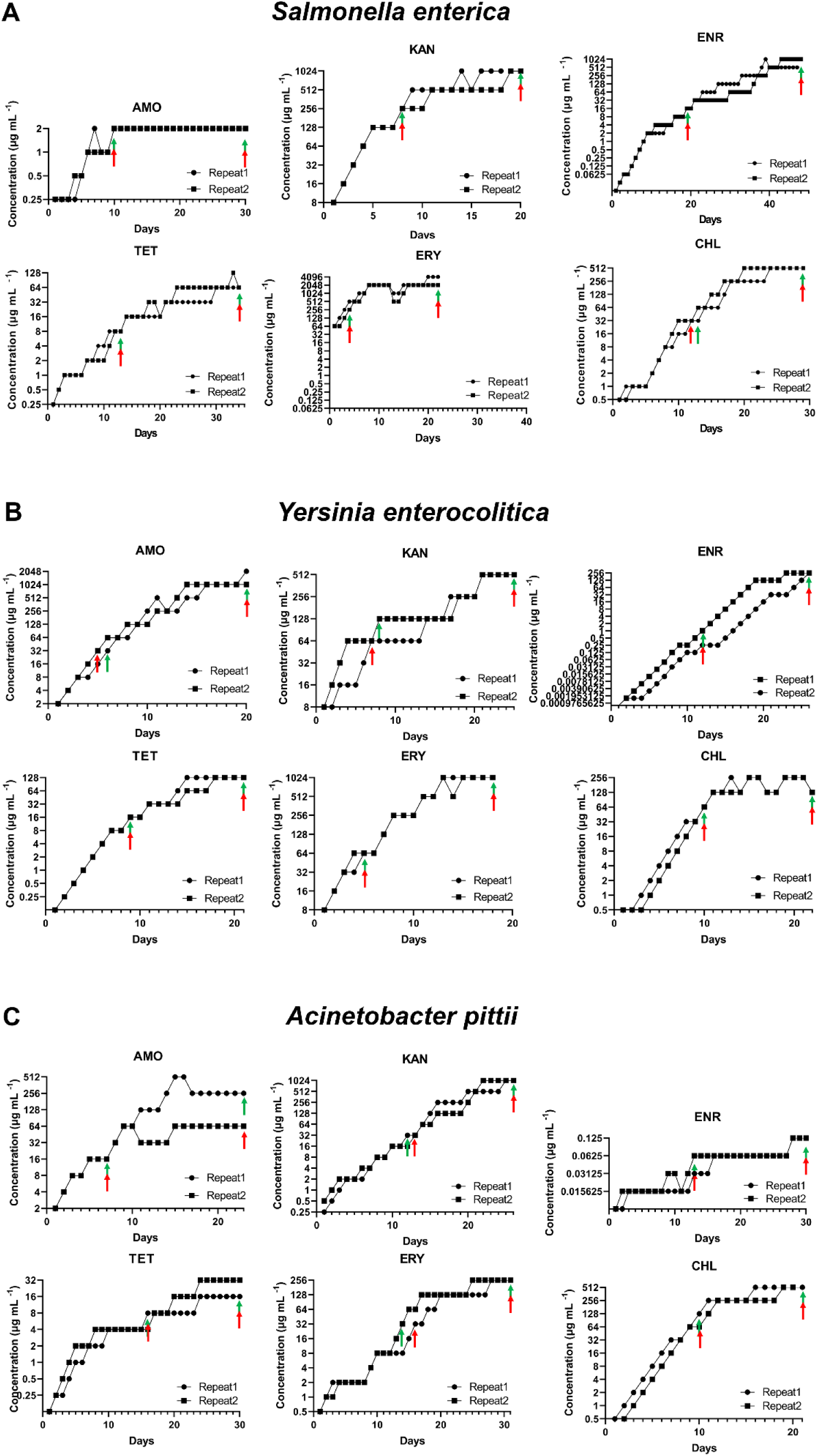
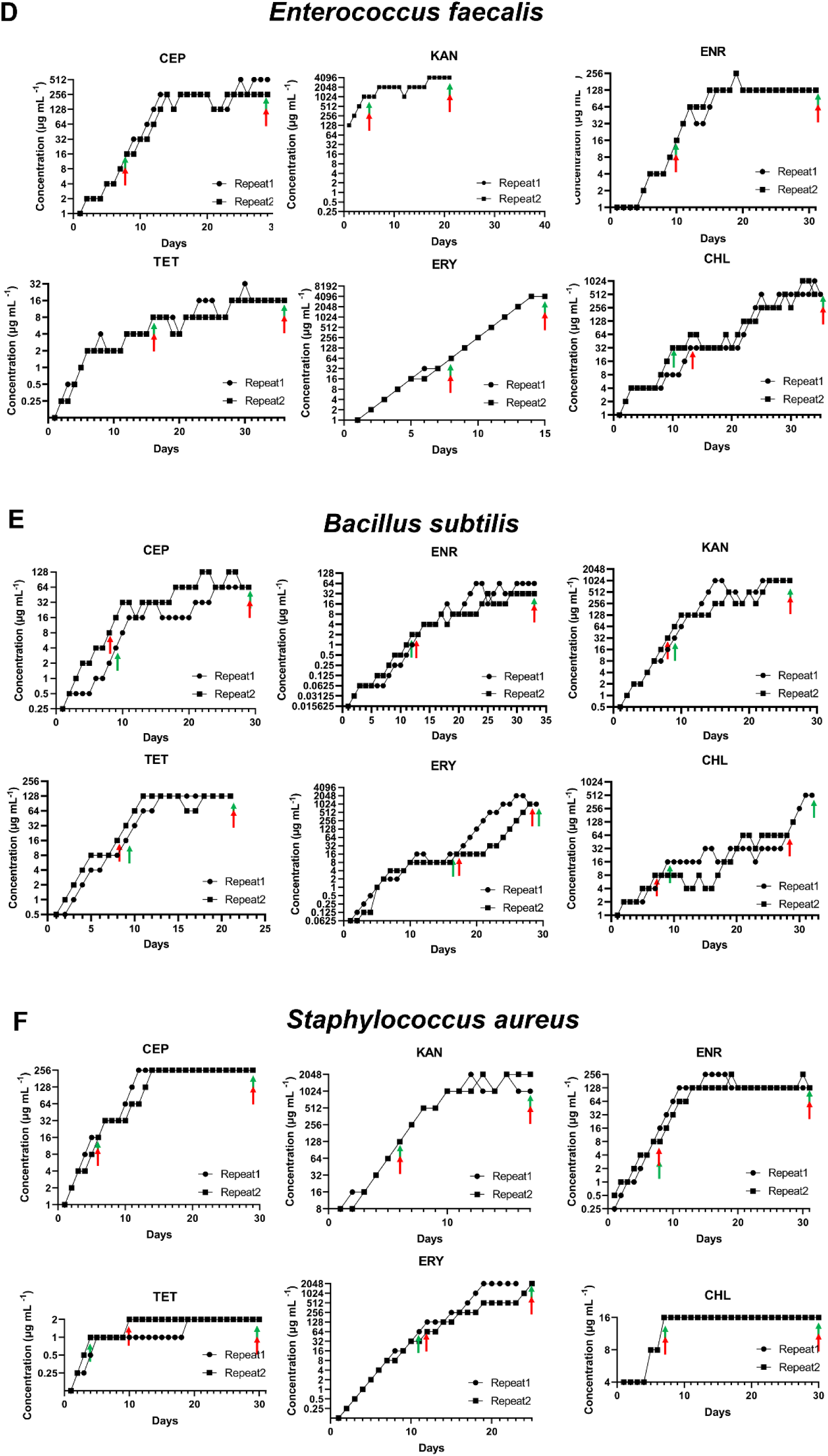
Resistance of six bacterial species evolved against six antibiotics. A) *S. enterica*; B) *Y. enterocolitica*; C) *A. pittii*; D) *E. faecalis*; E) *B. subtilis*; F) *S. aureus* The arrows indicate the days that samples for DNA analysis were taken; green indicates replicate 1 and red indicates replicate 2.

**FIG. 2.**
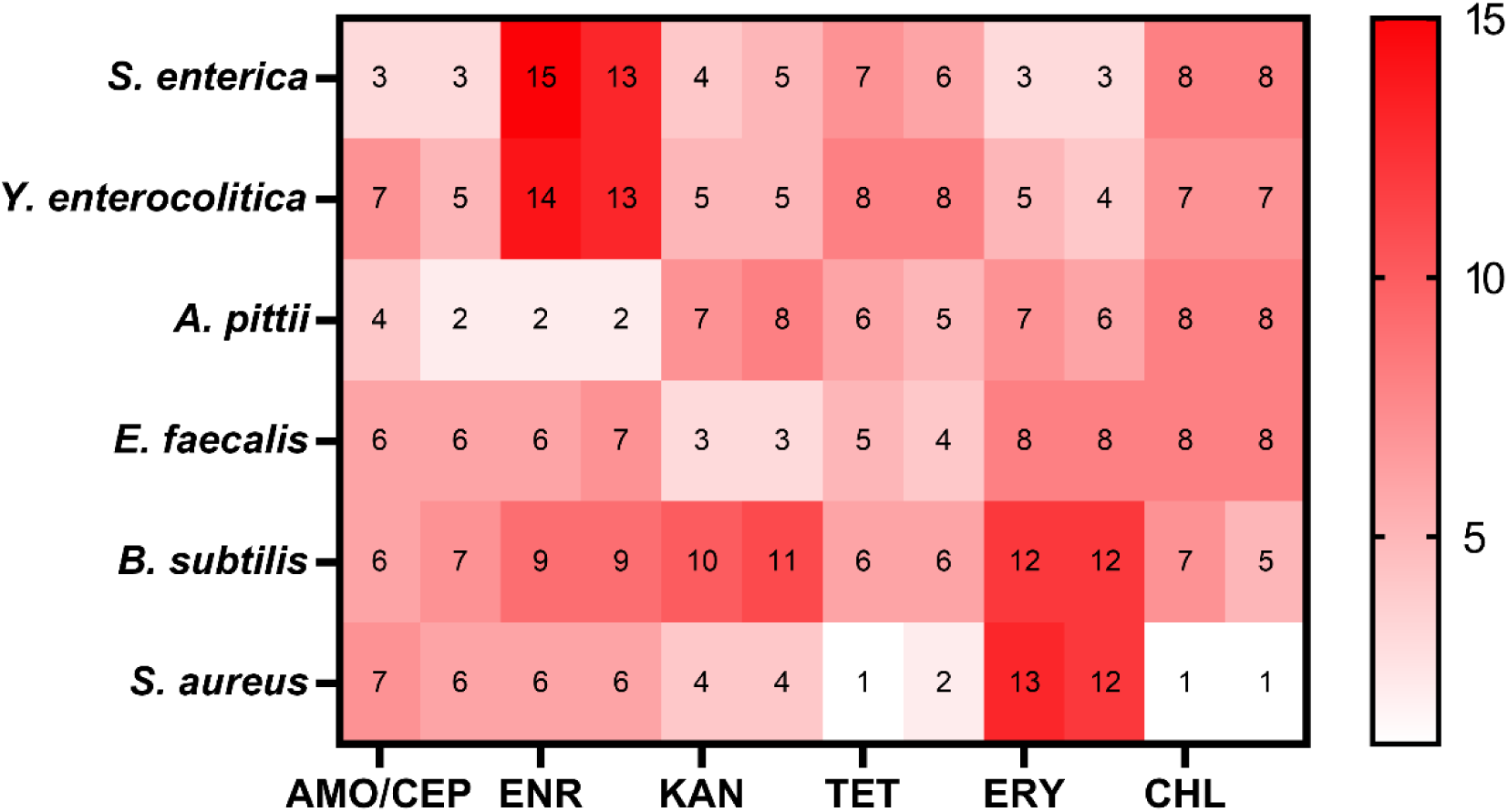
The MIC increase at the end of evolution in six species of bacteria. Six strains evolved resistance against amoxicillin/cefepime, enrofloxacin, kanamycin, tetracycline, erythromycin and chloramphenicol. The MIC of six strains at the end of the evolution experiment are compared to the MIC of the wildtypes. The MIC increase is expressed in 2^X^.

### Mutations associated with antibiotic resistance

To investigate the genetic mutations accompanying development of resistance during the evolution of antimicrobial resistance in these six strains, their DNA was sequenced at two time points (**Fig. 1**). The first measurement was roughly halfway the evolutionary trajectory, to document mutations contributing to initial resistance build-up. The second point at the end of evolutionary trajectory, aims to identify mutations that accompany the development of maximum resistance. All mutations in coding sequence regions and two promoter mutations of *rob* and *ampC* were categorized according to resistance mechanisms as defined in the CARD database and the related review (12). The six species had only a few mutations in common at the end of resistance evolution. Efflux pump related mutations were frequently encountered in gram-negative species. The effect of mutations on the phenotype, defined as MIC increase, is visualized in **Fig. 3**.

**FIG. 3.**
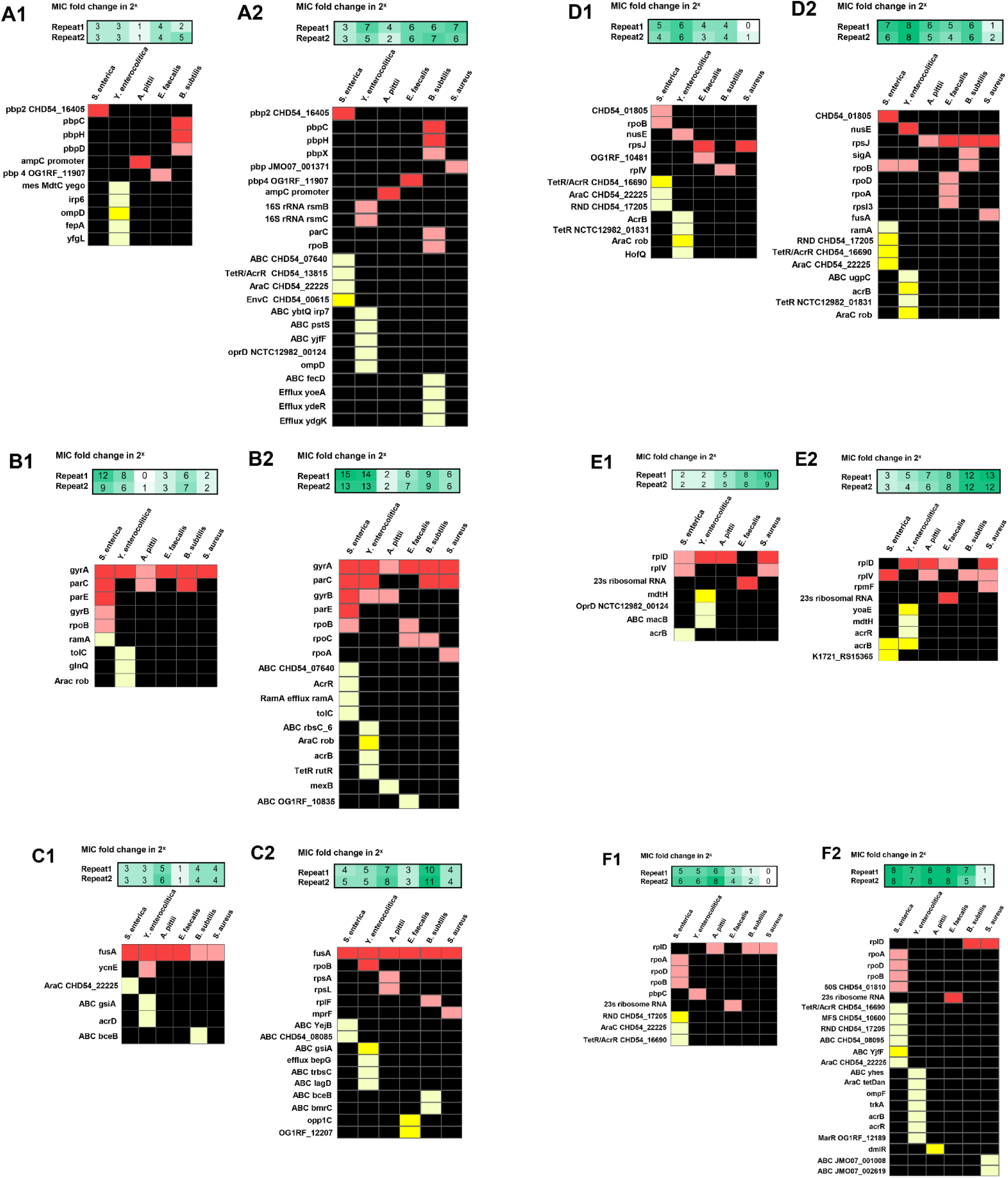
The distribution of point mutations in six species of bacteria. Mutations in early (1) or final (2) time point examined in A) amoxicillin/cefepime; B) enrofloxacin; C) kanamycin; D) tetracycline; E) erythromycin; F) chloramphenicol. Red color means target mutations; Yellow color means efflux pumps related mutations; Soft colors represent mutations found in one of the replicates; bright represents mutation found in both replicates.

Generally, DNA mutations involved in developing β-lactam resistance can be categorized into four main mechanisms: increased production of β-lactamase, modifications of penicillin-binding proteins (PBP), reductions of permeability, and increased efflux pump activity (20, 21). The gram-negative strains were evolved by exposure to amoxicillin and the three gram-positive strains evolved resistance against cefepime. The gram-negative *A. pittii* is the only strain that acquired a promoter mutation (-89 G>A) in *ampC*. Mutations in 16s rRNA *rsmB*/*rsmC* and mutation of genes related to efflux pumps were observed in *Y. enterocolitica*, which showed a more than 2^6^ increase in MIC to amoxicillin. In contrast, *A. pittii* and *S. enterica* developed little resistance to amoxicillin, as only a 2^3^ fold increase in MIC was observed. In *A. pittii* only mutations upstream of *ampC* (-89 G > A) occurred. *S. enterica* mutated *pbp2* and mutation of genes related to efflux pumps but did not develop significant amoxicillin resistance. The mutational pattern of the gram-positives was simpler than that of the gram-negative species. Only *B. subtilis* acquired mutations in a gene coding for efflux pumps. Mutations in *pbp4*, *pbpC*, *pbpX*, *pbpH* and *pbp JM007_001371* occurred during acquisition of cefepime resistance in *E. faecalis*, *B. subtilis* and *S. aureus*, accompanying around 2^6^ fold increase in MIC (**Fig. 3A**). Overall, the development of resistance to amoxicillin and cefepime was accompanied by mutations in *ampC* and *pbp*, respectively (**Table S1**). The number of known resistance mutations acquired during the full evolution is more than double as observed halfway through the evolution process (**Fig. 3A**).

During evolution of enrofloxacin resistance all six strains consistently mutated *gyrA*, in total, nine amino acid substitutions were observed in the *gyrA* locus. The serine around 83 was substituted in *S. enterica* (Ser83Phe), *Y. enterocolitica* (Ser83Ile), *E. faecalis* (Ser84Arg, Ser85Ile), *B. subtilis* (Ser84Leu) *A. pittii* (Ser81Leu) and *S. aureus* (Ser84Leu). The asparagine was the second target of amino acid substitution in *S. enterica* (Asp87Gly), *Y. enterocolitica* (Asp87Asn) and *A. pittii* (Asp70Gly). Additional replacement of glutamic acid was found in *E. faecalis* (Glu89Gly, Glu89Lys) and *B. subtilis* (Glu87Lys) (**Table S2**). In all strains, *parC* was mutated more often than genes coding for efflux pumps (**Fig. 3B1**). There were in total eleven amino acid substitutions in *parC*. The serine at 80 or 82 was the most frequently substituted amino acid: *S. enterica* (Ser80Phe, Ser80Arg), *B. subtilis* (Ser82Ile), and *S. aureus* (Ser80Phe) (**Table S3**). All evolved strains containing both *gyrA* and *parC* mutations had their MIC’s increased by more than 2^6^ fold, except for *A. pittii* (**Fig. 3B)**. A mutation in *parC* (Asn297Thr) with low allele frequency was observed halfway through the evolution in *A. pittii*, but it was lost during the subsequent adaptation process. To acquire the highest levels of resistance against enrofloxacin, bacteria combined several mechanisms. *Y. enterocolitica* and *S. enterica* had more than a 2^13^ fold increase in MIC, accompanied by mutations in *gyrA*, *parC* and genes coding for efflux pumps. The gram-positives *E. faecalis*, *B. subtilis* and *S. aureus* did not acquire resistance as much as the gram-negative species, despite their mutational patterns containing mutations in *gyrA*, *parC*, and the transcriptional regulators *rpoA*, *rpoB*, and *rpoC*. (**Fig. 3B2**).

Every strain made resistant to kanamycin contained a mutation in the gene coding for elongation factor G (*fusA*) (**Fig. 3C2**) In total, fourteen different amino acids were substituted in *fusA* across six species (**Table S3**). In gram-negative species mutations frequently occurred within the genes coding for the ATP-binding cassette (ABC) transporter permease and ABC transporter ATP-binding protein associated with an efflux pump. *B. subtilis* is the only gram-positive strain with mutations in the genes coding for the ABC family transporters (**Fig. 3C**).

In strains that evolved tetracycline resistance, mutations were found in genes coding for the 30S ribosomal protein, RNA polymerase and efflux pumps (**Fig. 3D**). Gram-negative bacteria developed higher resistance than gram-positives, and they acquired more mutations associated with the resistance–nodulation–division family (RND). More mutations in *rpoD* and *rpoB* were observed at the second time point (**Fig. 3D2**).

During development of erythromycin resistance, *Y. enterocolitica* mutated four genes coding for efflux pumps. *E. faecalis* acquired high level resistance, which is the only species that had mutations in 23S ribosomal RNA. *B. subtilis* did not have the well-known target mutations in genes for the 50S ribosomal protein, but still developed high level of resistance, accompanied by mutations of 5S, 16S, and 23S ribosomal RNA (**Fig. 3E**). In *S. aureus,* mutations were observed in *rplD*, *rplV* and *rpmF* and the MIC increased at least 2^12^ fold (**Fig. 3E2**).

The antibiotics chloramphenicol and erythromycin both target the 50S ribosome. Of the two, less target mutations were observed in the strains exposed to chloramphenicol. Similar to the other treatments, more mutations in genes coding for efflux pumps were documented in gram-negative strains (**Fig. 3F**).

### Species-specific response in antibiotic resistance evolution

To investigate the species-specific response as a result of antibiotic exposure, two replicates’ strains of six species were sequenced at the halfway and final point of their evolutionary trajectory (in total 144 measurements). The sequencing data were screened for mutations that accumulated in the bacterial genome and the dynamics of the allele frequencies were analyzed as well. Observed mutations were filtered using criteria (AF > 0.01 and Phred-quality score > 100). The number of DNA mutations differed widely between the six species. The highest number was found in *Y. enterocolitica* which accumulated more than 2,000 mutations at the end of the experiment. The other five species only accumulated a few hundred mutations (**Table 2**). A total of eight mutations in *mutS* or *mutL* were identified in *S. enterica*, *A. pittii* and *B. subtilis*. These strains acquired more mutations compared to other strains without mutator genotype (**Table S4**). In addition to that, the mutations of the two time points were dynamic as new mutations frequently replaced the initial ones. The majority of mutations are random, only a small proportion of early mutations later becomes dominant (**Fig. 4A**). At the end of evolution, six strains successfully acquired different levels of resistance against almost of antibiotics. However, the number of mutations that are fixed in the population (AF>0.9) differs in six species (**Table 2**). The mutational patterns in the six species were further analyzed based on mutation type and region (**Fig. 4B**). Single nucleotide polymorphisms (SNPs) were the most abundant mutation type in populations, with abundancies of 96.67% and 96.23% in *A. pittii* and *E. faecalis* relative to the total number of mutations. In the *Y. enterocolitica* strain, more short indels (<50 bp) were observed compared to the other species. Most mutations took place at the exonic regions. To determine the level of natural selection, the ratio of non-synonymous to synonymous (dN/dS) mutations was analyzed. The *B. subtilis* and *E. faecalis* were examined with the highest (4.512) and lowest selection (0.929), respectively.

**FIG. 4.**
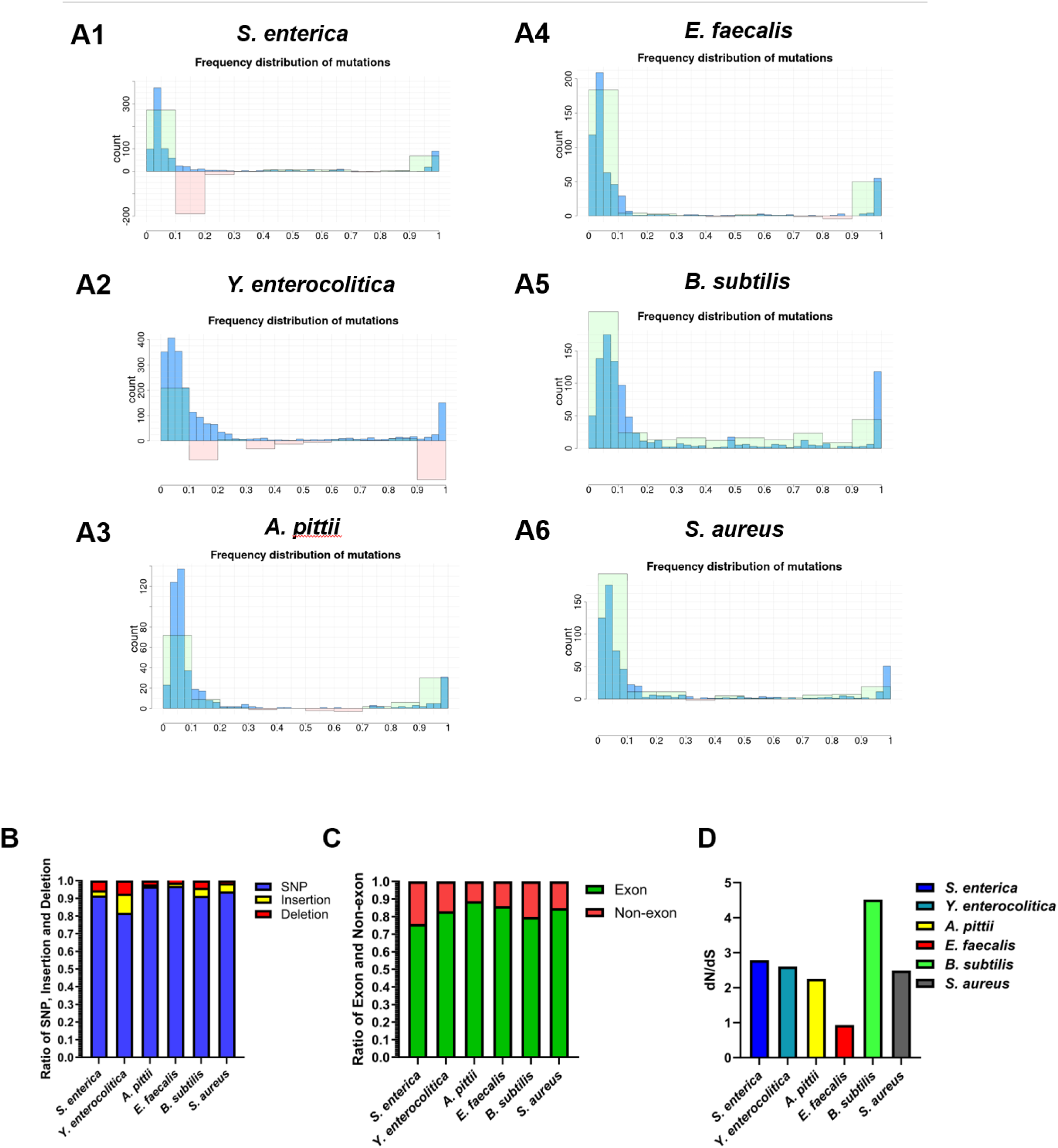
The distribution of allele frequency at the second measurement and analysis of mutation type. A1) *S. enterica*, A2) *Y. enterocolitica*, A3) *A. pittii*, A4) *E. faecalis*, A5) *B. subtilis*, A6) *S. aureus*; panels: the change in acquired mutations from measurement timepoint one to two is depicted with transparent green (positive change) and red (negative change) bars. B) The SNP, short insertion and deletion proportion. C) The proportion of mutations in the region of exon or non-exon. D) The ratio of non-synonymous mutations divided by synonymous mutations.

**TABLE 2.**
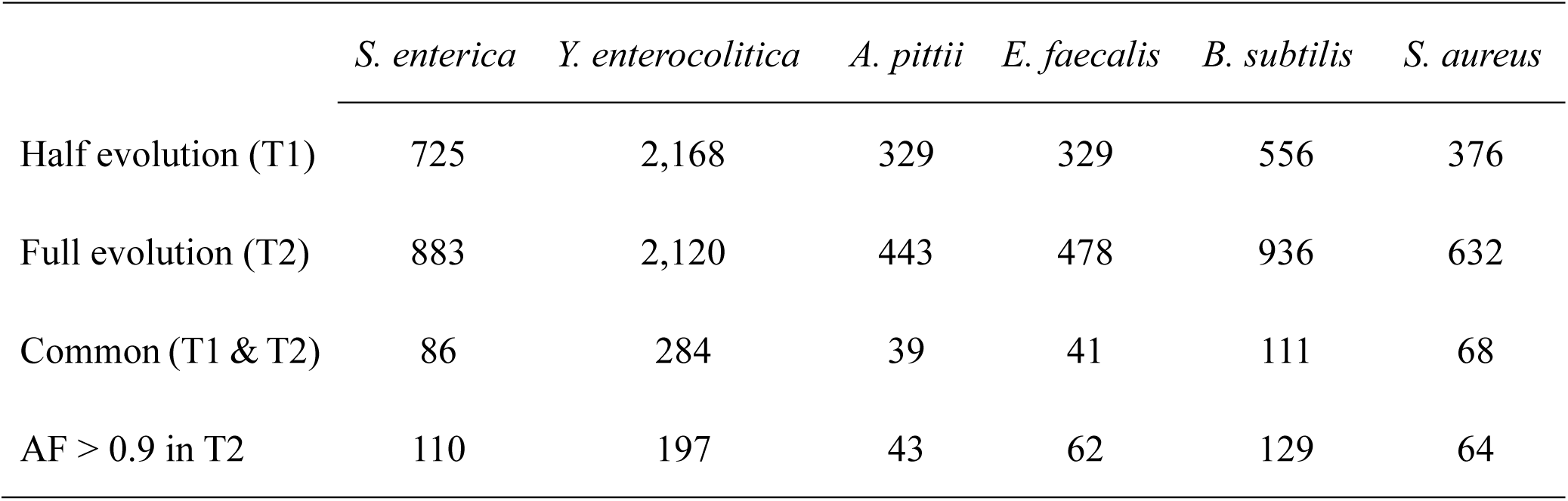
The number of mutations for each of the six bacterial species per timepoint.

### Impaired fitness as a consequence of resistance

To determine the metabolic trade-off of resistance, exponential growth rates were measured in wild type (WT) and resistant strains at two time points of evolution trajectory, as an indicator for fitness costs. Growth rates in *Y. enterocolitica* were significantly reduced during evolution compared to the other five species, accompanied by a higher frequency of SNP, insertion and deletion. (**Fig. 4B; Table 2**). *S. aureus* grew slower at the second measurement than the first measurement. In contrast to it, the other five species decrease similar degree of growth rate in these two measurements compared to wild type. Strains resistant to kanamycin had lower growth rates compared to the wild type (**Fig. 5**).

**FIG. 5.**
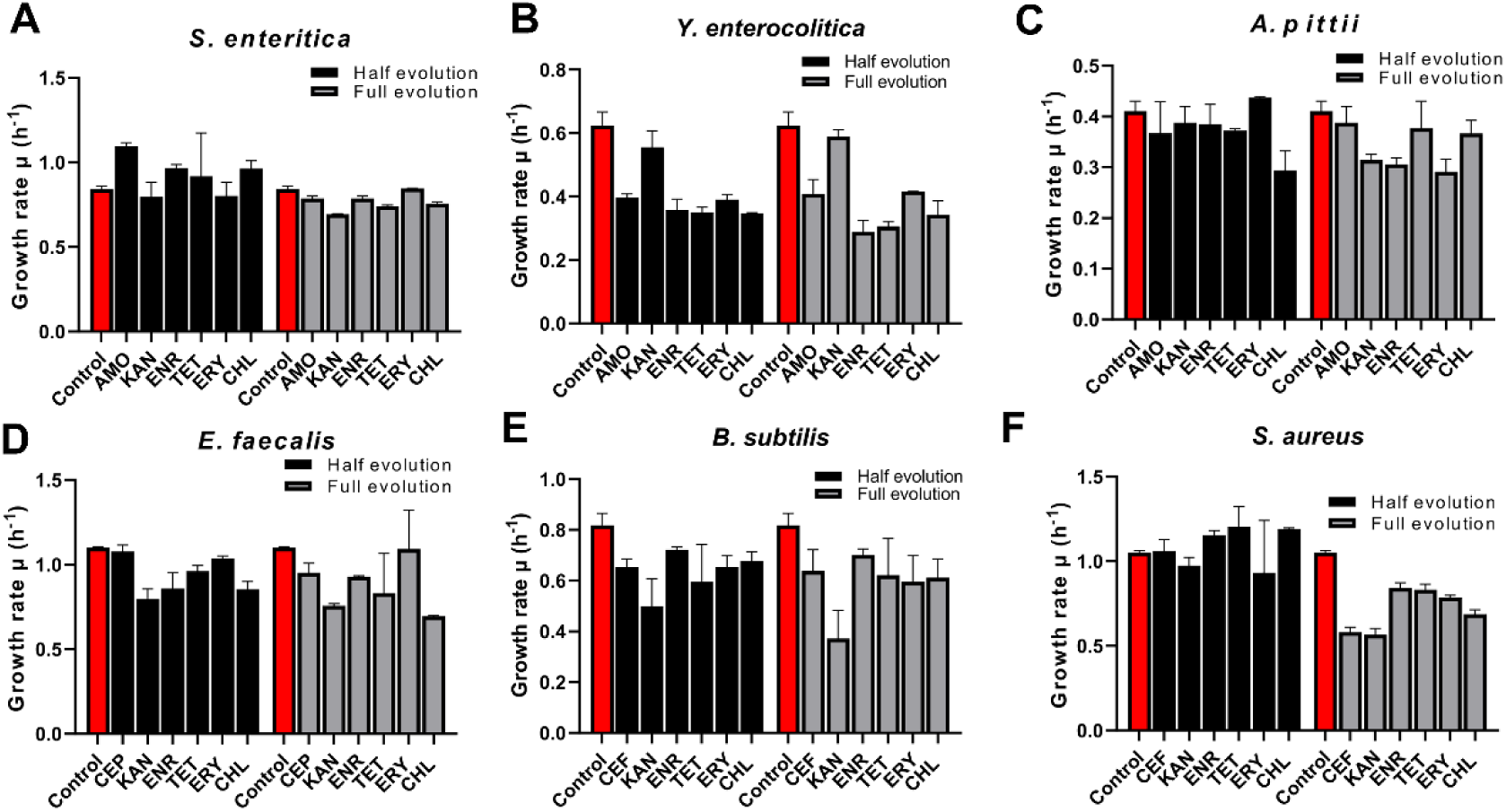
Growth rate as indicator for fitness cost measured in the absence of antibiotics of strains made resistant to the indicated antibiotic. A) S. enterica, B) *Y. enterocolitica*, C) *A. pittii*, D) *E. faecalis*, E) *B. subtilis*, F) *S. aureus*

**FIG. 6.**
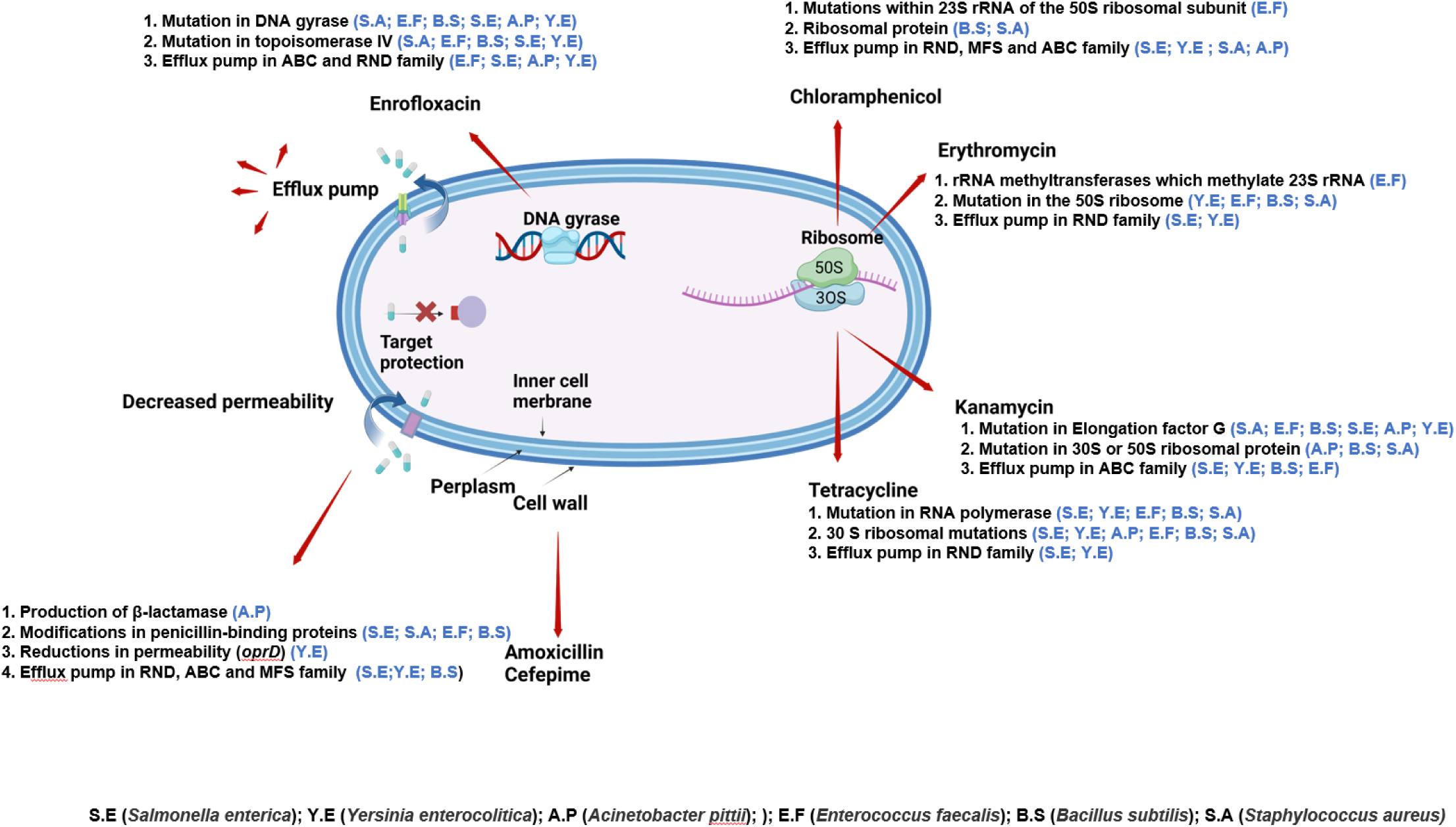
The mutational pattern are examined in six species of bacteria against six antibiotics.

## DISCUSSION

The evolution of *de novo* antimicrobial resistance has been extensively studied in *E. coli* (4–7, 22, 23), but more limitedly in other species such as *P. aeruginosa*, *A. baumannii and S. aureus* (24–26). Due to the variation in experimental conditions in these studies, it is difficult to make comparisons across species or antibiotics. Here, we investigated resistance evolution in six species relevant to veterinary microbiology to examine to which extent the conclusions based on development of resistance *E. coli* can be applied to more species. A second question addressed is whether there is a general pattern in the responses to specific antibiotics in these six species.

### Resistance

The evolutionary pathways towards resistance observed in the species investigated here were very similar to those in *E. coli* (4, 5, 7). No convergent evolutionary

### Mutations

The mechanisms involved in resistance against the antibiotics, amoxicillin/cefepime, enrofloxacin, kanamycin, tetracycline, erythromycin, and chloramphenicol include reduced permeability, antibiotic efflux, antibiotic modification, and target protection (11, 12). The promoter mutations in *ampC* found in *A. pittii* in this study were also reported in *E. coli* strains resistant to amoxicillin (7). The mutations in genes coding for PBPs that took place in *S. enterica*, *S. aureus*, *E, faecalis*, and *B. subtilis* in this study and were also reported in *P. aeruginosa* evolved against cefepime (17, 28, 29). The pair of mutations in *gyrA* and *parC* was found in *S. enterica*, *Y. enterica*, *A. pittii*, *B. subtilis* and *S. aureus* and reported in *K. pneumoniae*, *E. cloacae*, *A. baumannii*, *P. aeruginosa* and *S. pneumoniae* resistant to ciprofloxacin (13–15). The serine, glutamic acid and asparagine of *gyrA* and *parC* were the main amino acid targets for substitution in this study in strains evolved against enrofloxacin and the six strains of ESKAPE against ciprofloxacin (17). Similarly, the *fusA* is the most frequently mutated gene in strains exposed to kanamycin, consistent with the observations in *P. aeruginosa*, *E. cloacae*, *A. baumannii* and *K. pneumoniae* exposed to gentamicin (17, 24). Mutations in genes coding for efflux pumps are widely encountered in gram-negative bacteria. These mutations are likely to complicate treatment of infectious diseases (30, 31).

In bacteria evolving resistance against quinolones, clonal interference and epistasis influence rates of resistance development. The resistance level of populations is restricted by the fixation of mutations in *gyrA* and *parC* and mutations in genes coding for efflux pumps (13, 32). Usually, mutations in *gyrA* are the first step in the mutational trajectory of fluoroquinolone evolution. There is a competition between beneficial mutations of *gyrA* and mutations of *gyrB* in the early phase of ciprofloxacin evolution(13). However, the mutational pattern of bacteria evolved against enrofloxacin is different. We observed that mutations in *gyrB* took place after the mutations in *gyrA* in *S. enterica*, *Y. enterocolitica* and *A. pittii*, which is consistent with observations in *E. coli* (7). To acquire further resistance against fluoroquinolones, bacteria have to evolve additional mechanisms. The positive epistasis between mutations in *gyrA* and mutations in *parC* (7, 15), as well as mutations in *parC* and mutations in regulators of efflux pumps benefit strains by enabling them to adapt to substantial concentrations of fluroquinolone in *E. coli* (33). In this study, the resistance increased in all six strains when they acquired specific mutations in *parC* or beneficial mutations of efflux pumps, in agreement with observations in *E. coli* (7, 22), indicating that this type of sign epistasis is likely present in more species.

### Numbers of mutations

Bacteria have the ability to increase their mutation rate in reaction to antibiotic exposure (34). The large differences found in the number of mutations that accompany de novo resistance in *Y. enterocolitica* on the one hand and the other five species on the other, is consistent with the observations on more species. Like *Y. enterocolitica, E. coli* accumulated large numbers of mutations during resistance development (6, 22), while *Pseudomonas aeruginosa* had far fewer (14). Comparable high mutations rates were detected in *ampC*-depressed mutants of *Enterobacter cloacae* and *Enterobacter aerogenes* (35). Possibly the taxonomy reflects genetic characteristics, as *E. coli*, *E. cloaca, E. aerogenes* and *Y. enterocolitica* all belong to the order of *Enterobacterales*.

### Fitness cost

The fitness cost of antibiotic resistance development likely differs per species. Fitter resistant strains are likely to outcompete their resistant counterparts that contain DNA mutations (36). If however, the resistant strain has a lower specific glucose consumption, then the competitive edge is negligible and the extinction will not take place. In the agricultural sector in the Netherlands resistance levels started to decrease roughly two years after the initiation of a drastic reduction of the usage of antimicrobials(37). This indicates that the expected effect indeed occurs, but with a long hysteresis. The differences between the fitness costs incurred by the various species of this study can be explained in the framework Introduction of the hypothesis that fitness burdens of induced mutations depend on the genomic background (38). The overall fitness effect of resistance mutations is hard to predict, due to the epistasis and pleiotropy within the given genomic backgrounds (39)

### Mutational Dynamics during resistance development

The dynamics of DNA mutations during evolution towards resistance can be characterized as a trial-and-error process, as concluded for *E. coli* (7). As a consequence of exposure to non-lethal levels of antibiotics, the allele frequency of mutations with a positive fitness effect increases and the optimal mutations become dominant (13, 24). The frequency of crucial mutations for development of resistance rose to 100 % or close to that value. As part of this process new more beneficial mutations took over older mutations in population (40). Over all six species examined, only between 8 to 20 % of the mutations observed at the first time point were still present in the final measurement, again illustrating the dynamic and explorative nature of the process.

## LIMITATION

Due to a limited number of sequencing time points in the evolutionary trajectory, the dynamics of allele frequency of mutations is not fully examined and the change of diversity of mutations during the evolution is not completely documented.

## CONCLUSIONS

In this study, the six species can develop resistance for almost all of antibiotics. No common evolutionary trajectory was observed in the six species. Only a couple of mutations were commonly acquired in the six species, indicating that the species likely have their unique responses to antibiotic exposure. The genes *fusA*, *gryA* and *parC* are the most prevalent target of mutations. Mutation of genes coding for efflux pump were widely distributed in gram-negative species. *Y. enterocolitica* accumulated huge numbers of mutations and was observed the severe fitness cost in the process of building up resistance.

## ACKNOWLEDGMENTS

We appreciate that Sara Hernando Amado gave us constructive suggestions on earlier version of this manuscript. We thank Selina van Leeuwen for her assistance in DNA sequencing. The students Nimo Annor, Gioia Schaar and Sieradj Hendrikson performed experiments as part of their degree requirements. Xinyu Wang thanks the China Scholarship Council for supporting him through a PhD scholarship. This study was financed by The Netherlands Food and Consumer Product Safety Authority (NVWA). The sponsor was not involved in data collection or analysis, had no influence on the writing process and no permission was asked or given to publish this study.

## DATA AVAILABILITY STATEMENT

Most data underlying this article are available in the article and in its online supplementary material. Any further data are available from the corresponding author upon request.

## CONFLICTS OF INTEREST

The authors declare that no conflict of interests, financial or otherwise, exist for any of the authors.

